# Longitudinal monitoring of disease burden and response using ctDNA from dried blood spots in xenograft models

**DOI:** 10.1101/2022.01.15.476442

**Authors:** Carolin M Sauer, Katrin Heider, Jelena Belic, Samantha E Boyle, James A Hall, Dominique-Laurent Couturier, Angela An, Aadhitthya Vijayaraghavan, Marika AV Reinius, Karen Hosking, Maria Vias, Nitzan Rosenfeld, James D Brenton

## Abstract

Whole genome sequencing (WGS) of circulating tumour DNA (ctDNA) is now a clinically important biomarker for predicting therapy response, disease burden and disease progression. However, the translation of ctDNA monitoring into vital pre-clinical PDX models has not been possible owing to low circulating blood volumes in small rodents. Here, we describe the longitudinal detection and monitoring of ctDNA from minute volumes of blood in PDX mice. We developed a xenograft Tumour Fraction (xTF) metric using shallow WGS of dried blood spots (DBS), and demonstrate its application to quantify disease burden, monitor treatment response and predict disease outcome in a pre-clinical study of PDX mice. Further, we show how our DBS-based ctDNA assay can be used to detect gene-specific copy number changes and examine the copy number landscape over time. Use of sequential DBS ctDNA assays will transform future trial designs in both mice and patients.

## Introduction

Liquid biopsies are routinely used in the clinic to sensitively detect and quantify disease burden, and have critical roles for therapeutic decision making in precision medicine^1–6^. Plasma circulating tumour DNA (ctDNA) is the most widely studied circulating analyte for disease monitoring and molecular genotyping of tumours^2,7^. Technical advances in next generation sequencing (NGS) now achieve unprecedented sensitivities for the detection of ctDNA using 6-10ml of whole blood^1,8^. To enable very accurate monitoring of disease burden and progression, several whole genome sequencing (WGS)-based strategies have been developed detecting combinations of single-nucleotide variants, small insertions/deletions and somatic copy number aberrations (SCNAs)^9–14^. In addition, deriving other biochemical features of ctDNA from WGS, including fragment size and chromosome accessibility, can further enhance detection sensitivity and infer biological information about tumour site of origin^15–19^.

Modelling therapeutic response in mice bearing patient-derived xenografts (PDX) is a critical step to test treatment regimens and pharmacogenomics during drug development^20,21^. However, WGS-based ctDNA assays cannot be used in small rodents as the circulating blood volume of a mouse is only ~1.5-2.5ml. Consequently, detailed ctDNA assays can only be obtained from terminal bleeding of mice, preventing longitudinal analyses and more efficient therapeutic study designs. Manual measurements of tumour volumes in subcutaneous models are the commonest surrogate to estimate treatment response and disease burden^22,23^. These measures are often poorly reproducible and can be biased by treatment-induced tissue necrosis and oedema. Using imaging as an alternative to estimate response in PDXs is more time-consuming, requires general anaesthesia and may also need the introduction of *in-vivo* reporter genes^24,25^.

Therefore, bringing WGS-based ctDNA assays into mice would have two major benefits: firstly, more efficient and accurate serial measurements across multiple animals, and secondly the direct translation of biological and biochemical observations from mouse ctDNA studies into patient studies and vice versa. We recently illustrated the detection of ctDNA in dried blood spots (DBS) from minute volumes of whole blood using a size selection approach to enrich for cell free DNA (cfDNA)^26^. Using a modified approach in PDX mice, we now demonstrate that shallow WGS (sWGS) of DBS from 50μl of whole blood can be used for serial ctDNA measurements, longitudinal disease monitoring and copy number analyses in pre-clinical studies. The work presented here provides important proof-of-principle data and further supports the application and feasibility of DBS-based ctDNA sampling both in pre-clinical and clinical studies.

### Development and validation of the xTF metric from DBS

To detect and accurately quantify ctDNA from minute volumes of blood in pre-clinical PDX studies, we developed a xenograft Tumour Fraction (xTF) metric, which is estimated from shallow whole genome sequencing (sWGS) of DBS samples (**Fig. 1a**). Briefly, 50μl of blood are collected from the tail vein, deposited onto a filter card and left to air-dry. DNA is extracted, contaminating genomic DNA is removed^26^ and subsequently sequenced at low coverage following library preparation. Human- and mouse-specific reads are identified using Xenomapper^27^, and the xTF is calculated as the ratio of human-specific reads divided by total reads (human and mouse specific reads) per sample (**see Methods**).

**Figure 1 –.**
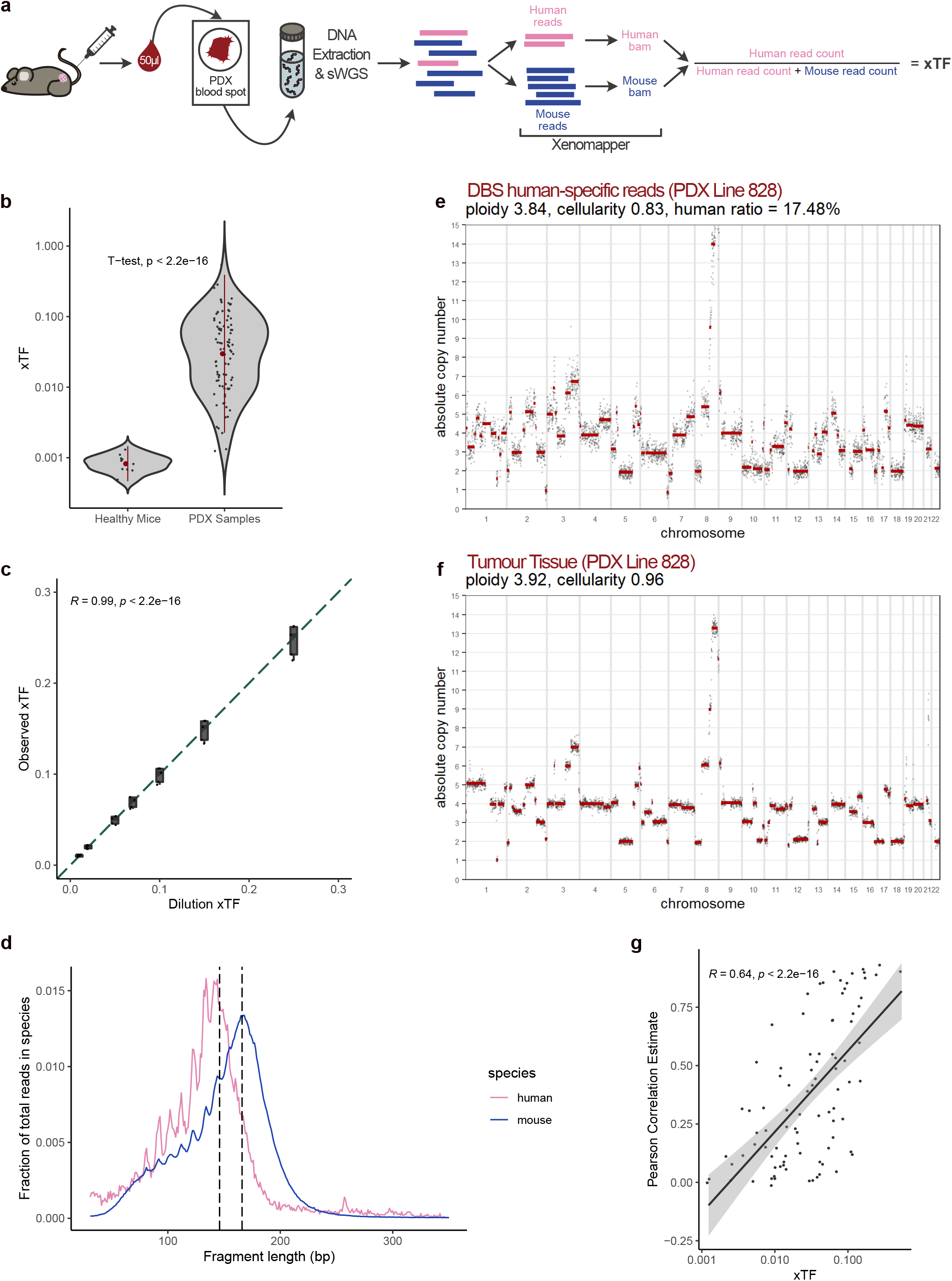
The xTF metric is highly specific and sensitive to detect and quantify ctDNA from dried blood spots. (**a**) Workflow of the dried blood spot (DBS)-based xenograft Tumour Fraction (xTF). DBS are generated by collecting and depositing 50 μl of blood from the tail vein of the mouse onto FTA filter cards. DNA is extracted from blood spots, processed and sequenced as described previously^26^. Human-specific reads and mouse-specific reads were separated into species-specific bam files using Xenomapper^27^. The xTF is then calculated by dividing the number of human specific reads by the total number of human and mouse specific reads in a given sample. (**b**) Comparison of xTF values obtained from healthy non-tumour bearing mice and PDX samples (Welch t-test, p < 2.2 x10^-16^). Sensitivity testing using the Mann-Whitney *U* Wilcoxon test (Wilcoxon test, p = 2.5 x10^-7^) showed similar results. (**c**) xTF dilution series. Dilution xTFs (0.01, 0.02, 0.05, 0.07, 0.1, 0.15 and 0.25) were computationally generated by mixing blood spot sequencing data obtained from five ovarian cancer patients and a healthy control mouse. The generated dilution series was analysed using Xenomapper and resulting xTF values were compared to the dilution xTFs (Spearman correlation R = 0.99, p < 2.2 x10^-16^). (**d**) Fragment length distributions of human- (pink) and mouse- (blue) specific reads from a DBS sample. Two vertical lines indicate 146 and 166 bp, the observed peaks for ctDNA and cfDNA, respectively. (**e**) Example of an absolute copy number (ACN) profile successfully generated from human-specific reads from a DBS collected from a PDX mouse of patient line 828. (**f**) Matching ACN profile generated from sWGS of PDX tumour tissue. **Supplementary Fig. 1 and 2** show representative ACN profiles for all four patient lines and the correlation of each copy number bin for the DBS and tissue sample pairs. (**g**) Correlation of Pearson correlation estimates (comparing ACN bins between tumour tissue and DBS) and xTFs from DBS samples (Spearman R = 0.64, p < 2.2 x10^-16^).

To test both the specificity and sensitivity of the xTF metric, we established a pre-clinical study using PDX mice derived from four high grade serous ovarian cancer (HGSOC) patients (see next section). We collected a total of 10 DBS samples from 5 healthy non-tumour bearing mice and 91 DBS samples from 35 tumour bearing PDX mice. Reads from healthy control mice showed <0.1% assignment as human-specific sequences (false-positive background). In addition, healthy control mice had significantly lower xTF values compared to tumour-bearing PDX mice, independent of tumour size and disease burden, indicating the high specificity of the xTF metric (Welch t-test, P = 2.2×10^-16^, **Fig. 1b**). To confirm the linearity and sensitivity of our approach, we prepared an *in silico* 7-point dilution series (**see Methods**) by combining sequencing reads from a healthy mouse DBS and DBS samples collected from five independent ovarian cancer patients at different ratios. We were able to accurately detect human reads for all seven dilution points, and observed a strong correlation between measured xTFs and spiked-in human reads at human:mouse proportions of 1-25% (Spearman’s R = 0.99, P < 2.2×10^-16^, **Fig. 1c**)

Next, we examined the fragment size distributions of human- and mouse-specific reads from sWGS of DBS samples. In human plasma samples, ctDNA has a modal size of approximately 145bp, which is shorter than cfDNA with a prominent mode of approximately 165bp^15,28^. These fragment size properties were recapitulated in the human- and mouse-specific reads from DBS samples (**Fig. 1d**). Given the high specificity and sensitivity of our approach, we were able to derive absolute copy number (ACN) data from as little as 500,000 human-specific DBS reads using QDNAseq^29^ followed by Rascal^30^ (**Fig. 1e**). The observed absolute somatic copy number aberrations (SCNAs) (**Fig. 1f**) and their extent were strongly correlated with sWGS of PDX tumour tissues from the same patient (**Supplementary Fig. 1 and 2a-d**). Unsurprisingly, the ability to accurately detect SCNAs in DBS strongly correlated with increasing xTF values (**Fig. 1g**). No correlations were observed when comparing blood spot ACN profiles from healthy non-tumour bearing mice to any of the four patient tumour tissues (**Supplementary Fig. 2e-g**). Using the definitions of copy number gains and losses outlined by the Catalogue of Somatic Mutations In Cancer (COSMIC), amplifications of driver SCNAs were detectable in blood spot samples with xTFs ranging from 0.6-54.4% (**Supplementary Fig. 1 and 2h**).

### The xTF allows accurate monitoring of disease progression

We next investigated whether the DBS-based xTF assay could be used for longitudinal monitoring of disease progression and treatment response. An overview of our pre-clinical PDX study is shown in **Fig. 2a**. The PDX models were selected from 4 patients with different clinical responses to platinum-based chemotherapy and distinct copy number signatures^31^ for homologous recombination deficiency (HRD) that are predictive of sensitivity to carboplatin (**Supplementary Fig. 3 and 4**). All PDXs were derived from tumour samples prior to systemic therapy and histological and molecular features were shown to be highly similar to the primary tumour (**Supplementary Fig. 5 and 6**). PDX mice were treated with either 50mg/kg carboplatin or control on day 1 and 8. Tumour volumes were measured weekly, and blood spots were collected on day 1 (prior to treatment start), day 16 and 29 (**Fig. 2a**).

**Figure 2 –.**
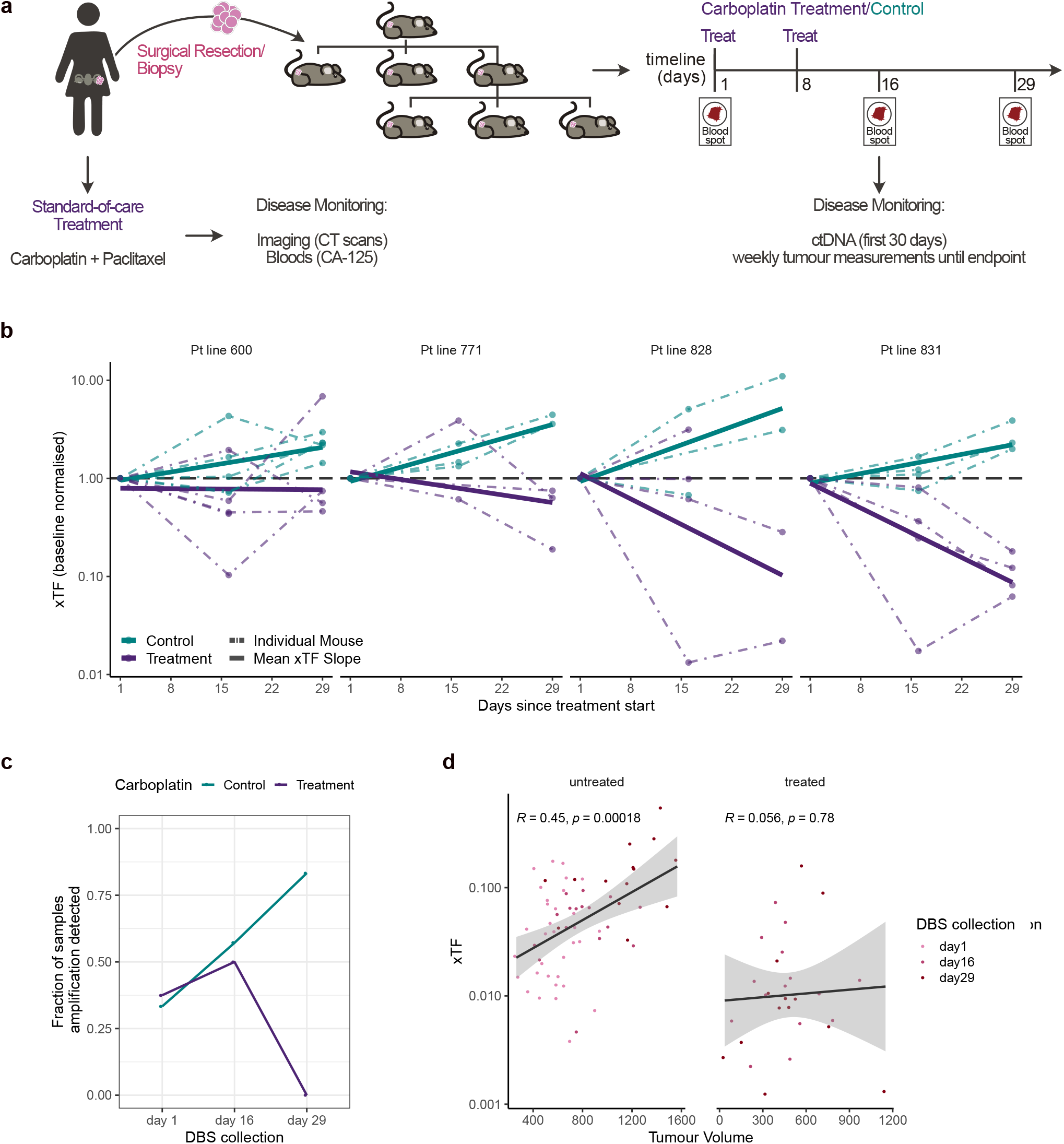
The DBS-based xTF allows longitudinal monitoring of disease progression and treatment response in pre-clinical studies. (**a**) Pre-clinical PDX study overview. HGSOC patients underwent surgery and standard-of-care chemotherapy with carboplatin and paclitaxel. Disease progression was monitored over time using the CA-125 biomarker, CT scans, as well as ctDNA where available. The treatment-naïve surgical tumour or biopsy specimens were engrafted into NSG mice. Second or third generation PDX mice were then treated with either carboplatin or vehicle control via tail vein injection on day 1 and day 8. Tumour volumes were measured weekly, and blood spots were collected on day 1 (prior to treatment initiation), day 16 and 29. (**b**) xTF change from baseline during the first 29 days since treatment start for each PDX patient line. xTFs were normalised to baseline (day 1) xTF values for each mouse (dashed lines). Carboplatin-treated mice are shown in purple, control mice are shown in teal. Bold lines show the linear-model fitted line across all mice within the same treatment and patient group. Horizontal dashed lines at y=1 indicate normalised baseline. (**c**) Fraction of blood spot samples in which putative driver amplifications were detected over time. The fraction of samples with detected gene amplifications decreases in carboplatin treated group, while increasing in the control group over time. (**d**) xTF values were correlated with tumour volumes of the nearest matched time-point for both untreated (Spearman R = 0.45, p = 0.00018), and carboplatin-treated (Spearman R = 0.056, p = 0.78) PDX mice.

We observed a progressive increase in xTF in all 17 untreated PDX control mice. In contrast, the 18 mice that were treated with carboplatin showed variable decreases in xTF detected in DBS samples collected at day 16 and 29 in comparison to pre-treatment (day 1) samples (**Fig. 2b**). Similarly, the fraction of samples in which we were able to detect human gene-level amplifications (i.e. *MYC* and *MCM10* amplifications in patients 828 and 771, respectively) from DBS reads increased in untreated and decreased in carboplatin-treated mice over time (**Fig. 2c**). When correlating xTF values to tumour volumes obtained from weekly tumour measurements, we found that xTFs increased with increasing tumour volumes and thus disease burden in untreated control mice (Pearson’s R = 0.45, P = 0.00018, **Fig. 2d**). However, no correlation was found in treated samples (Pearson’s R = 0.056, P = 0.78, **Fig. 2d**), likely because of treatment-induced tissue necrosis and oedema biasing tumour volume measures.

### The xTF rate of change is predictive of disease outcome

Early dynamic change in ctDNA can predict progression-free survival and provide real-time assessment of treatment efficacy^32^. Similar predictive measures in mice could also improve the efficiency of PDX study designs. All four PDX lines in our cohort were from patients with platinum-sensitive disease, and PDX 828 and 831 were predicted to have the best response to carboplatin treatment owing to somatic and germline *BRCA1* mutations, respectively (**Supplementary Fig. 3 and 7**). PDX 600 and 771 had less marked HRD signatures (**Supplementary Fig. 4 and 7**). Clinical progression free survival (PFS) and overall survival (OS) (**Supplementary Fig. 7**) could not be used as response predictors as the four patients have important differences in prognostic variables for stage and residual disease after surgery (**Supplementary Fig. 3**).

We asked whether the rate of change in xTFs during the first 30 days following initiation of treatment was predictive of disease outcome in our PDX cohort. Given the poor correlation between xTFs and tumour volumes (**Fig. 2d**), we explored tumour growth kinetics from weekly tumour measurements taken from the time of tumour engraftment until study endpoint for carboplatin-treated and untreated mice (**see Methods**, **Fig. 3a-d**). Tumour volumes and growth rates were not significantly different between treatment and control mice across the four lines prior to start of treatment (**Supplementary Table 1**). Importantly, the rates of tumour regrowth in treated mice were not significantly different from initial growth rates after engrafting and prior to treatment start (**Supplementary Table 1**), providing evidence that carboplatin treatment (and potential clonal selection) did not change tumour growth kinetics. We then inferred inflection points representing treatment-induced changes in tumour growth rates, allowing estimation of both the time of treatment response (t_1_) and time of tumour regrowth (t_2_) (**Fig. 3a-d**). t_2_-t_1_ therefore represents the duration of treatment effect, and t_2_ is comparable to PFS, the commonest clinically validated surrogate endpoint for clinical trials. As predicted, t_2_-t_1_ measures were longest (best response) for PDX 828 and 831 and the worst response was seen in PDX 600.

**Figure 3 –.**
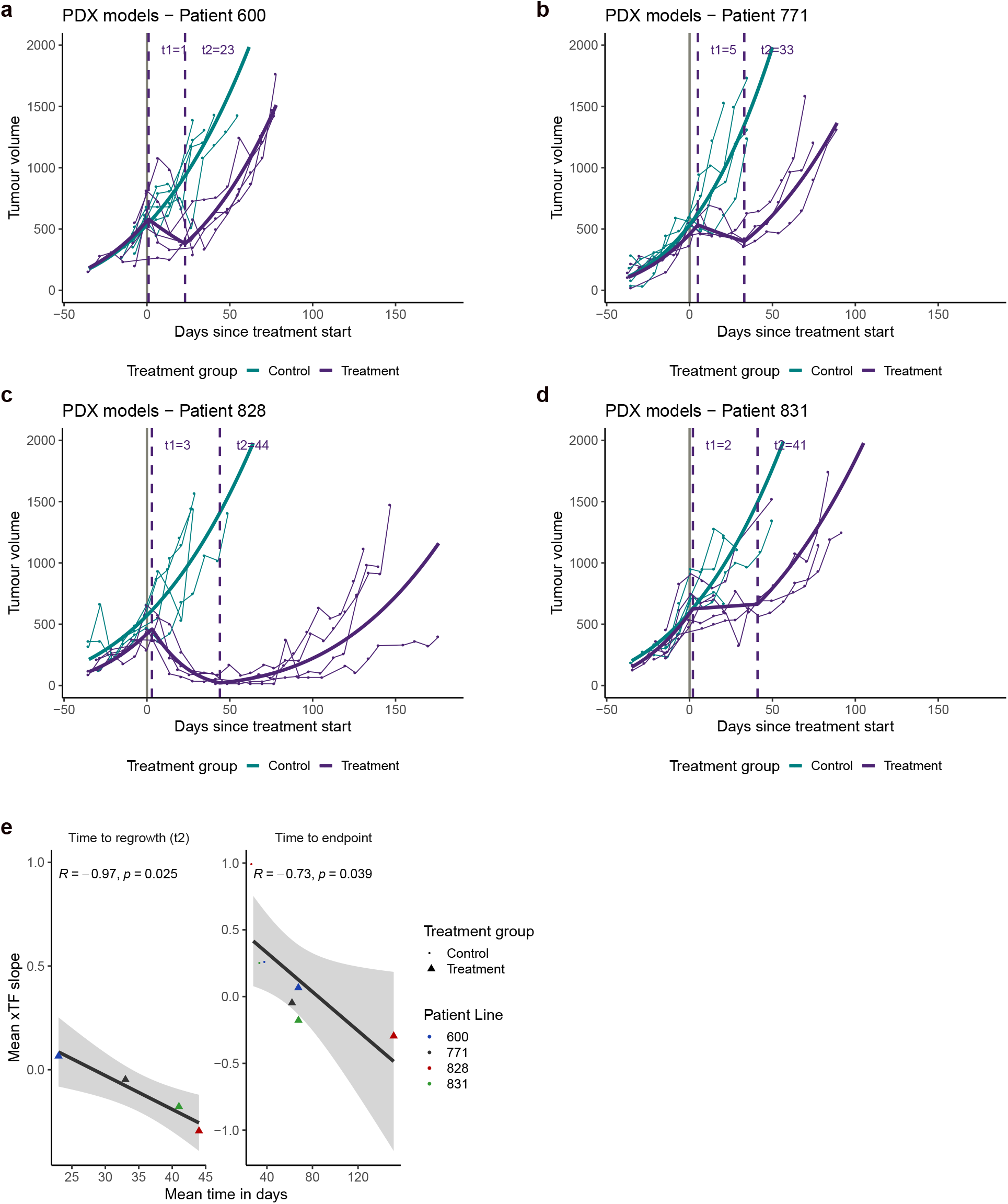
Change in xTF over time predicts disease outcome. (**a-d**) Weekly measured tumour volumes (mm^3^) for PDX mice over time. Treatment start is indicated by grey solid vertical line. Solid coloured lines show modelled tumour growth curves using heteroscedastic point-wise random intercept linear mixed models (**see Methods**). Growth curve inflection points were determined (**see Methods**) to estimate the start of treatment effect and tumour regrowth (dashed vertical purple lines labelled t_1_ and t_2_, respectively). (**e**) The mean xTF slope was estimated for each treatment group across the four patient lines (**Fig. 2b**) and compared to the mean time to endpoint (Spearman R = −0.73, p = 0.039) and tumour regrowth (Spearman R = −0.97, p = 0.025). The four different patient lines are indicated by different colours.

Importantly, there was a strong negative correlation between xTF change rate (**Fig. 2b**) during the first 30 days of treatment and t_2_ (tumour regrowth; Pearson’s R = −0.97, P = 0.025). xTF was also strongly correlated with study endpoint (a surrogate for overall survival; Pearson’s R = −0.73, P = 0.039) (**Fig. 3e**).

## Discussion

We here demonstrate for the first time how minimally-invasive sampling of DBS can be used to accurately monitor disease progression and treatment response in PDX mice using sWGS of ctDNA. The low volume of blood required allows repeated serial collection of ctDNA samples from living, non-anaesthetized mice and removes the need for terminal bleeding. Further, detailed modelling of tumour response indicates that the initial change in xTF in response to treatment is predictive of PFS and OS. This advocates the use of the xTF metric as a reliable minimally-invasive tool to monitor disease progression and to study treatment response in pre-clinical settings.

DBS are derived from whole blood; sensitive detection of ctDNA from DBS therefore requires removal of contaminating genomic DNA which otherwise significantly dilutes ctDNA signal^26^. In comparison to plasma samples, however, DBS have clear sampling advantages, since they do not require prompt centrifugation, and provide stable and space-efficient storage of DNA for many years^33^. DBS therefore have the potential to simplify sample collection and revolutionise study designs in both mice and patients:

In mice, the use of DBS has already been illustrated in pharmacokinetic studies^38^ and has proven to conform with the 3Rs of animal welfare^39^ by reducing the number of animals required per study, allowing facile sample collection at multiple timepoints, and improving the quality and quantity of data collected from a given mouse. In this study, blood samples were collected from the tail vein, which is considered a simple, humane and anaesthesia-free approach^40^. Alternative methods include submandibular or saphenous bleeding^41,42^ which, in contrast to tail vein bleeding, do not require the use of a mouse restrainer and will preserve the tail vein for drug administration. Although tail vein blood sampling has previously been optimised for disease monitoring, ctDNA assays were limited to PCR-based experiments from plasma^43^. In contrast, sWGS of DBS can be used to simultaneously assay ctDNA features and the copy number landscape of engrafted tumours. We show that mouse- and human-specific reads recapitulated the fragment size properties of human cfDNA and ctDNA, respectively, indicating that mechanisms of cfDNA/ctDNA release into the blood stream is similar in mice and humans. Our approach is therefore a promising platform to study factors influencing ctDNA shedding, as well as other biochemical features of ctDNA, such as methylation and nucleosome profiles.

In the clinic, DBS-based technologies may allow self-collection at home (via a simple finger-prick), obviating the need for additional phlebotomy or hospital visits, and thus improving test acceptability and study participation. While our approach proved to be highly sensitive for the detection and quantification of disease in PDX mice, it relied on the ability to identify tumour-specific (human) ctDNA reads from DBS sequencing data using species-specific read alignment. This will not be possible in DBS samples collected from cancer patients. However, similar sensitivities might be achieved by implementing fragmentomic^15,18^ or epigenomic features^34^ and patient-specific mutations (personalised sequencing panels)^10,11,35–37^ for ctDNA detection, facilitating sensitive disease monitoring from small blood volumes in the clinic^17^.

In summary, we reported the unprecedented use of WGS of ctDNA in murine models, which provides a powerful new tool for pre-clinical disease monitoring and allows accurate assessment of treatment response and simultaneous assaying of the copy number landscape over time. This provides an exciting opportunity for future research to study copy number driven tumour evolution and to investigate how treatment-induced selection of copy number changes may result in treatment resistance. Importantly, the use of DBS-based ctDNA assays provides an important opportunity to simplify and improve study design in both mice and patients.

## Methods

### Generation of PDX mouse models

Solid tumour samples were obtained from patients enrolled in the OV04 study (CTCROV04) at Addenbrooke’s Hospital, Cambridge. Tumour samples were processed following standardised operating protocols as outlined in the OV04 study design and as previously described^30^ before surgically engrafting into female NOD.Cg-Prkdc^scid^ Il2rg^tm1WjI^/ SzJ (NSG) mice obtained from Charles River Laboratories. All mouse work conducted was approved and performed within the guidelines of the Home Office UK and the CRUK CI Animal Welfare and Ethics Review Board. Xenograft tissue processing and PDX passaging were performed as previously described^30^. In short, xenografting was performed either by subcutaneous surgical implantation (for first generation PDX mice) or subcutaneous injection of tumour cells from dissociated tumour tissues (for later PDX generations). Tumour bearing mice that reached their endpoint (tumour volumes of no more than 1500mm^3^) were culled via cervical dislocation or CO_2_ overexposure. Tumour tissues were dissected, processed as described above and re-transplanted for expansion in serial generations for PDX biobank maintenance and model generation.

### Treatment of mice

Treatment was initiated when engrafted tumours reached a size of approximately 500mm^3^. Mice were randomised to either receive 50mg/kg of carboplatin (dissolved in water for injections (WFI) and mannitol (10mg/ul)), or 100μl carboplatin vehicle/control (10mg/ml of WFI diluted mannitol).

Mice were treated by tail vein injection on day 1 and day 8 and monitored until they reached their endpoint of 1500mm tumour volume, or if another health concern was raised.

### Measurement of tumour volume

Using callipers, the height (*h*), width (*w*) and depth (*d*) of the mouse tumours were measured in millimetres once a week and the tumour volume (mm^3^) was determined using the formula:

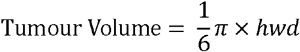

### PDX tumour growth curve modelling

Heteroscedastic point-wise random intercept linear mixed models were used to model the tumour growth (on the cube root scale) of both, control and treated mice for each of the four patients included in this study. Heteroscedastic models were preferred as the (tumour growth) variance of treated mice appeared larger than the variance observed in the control mice.

For carboplatin treated mice within each patient group, the time points of the following two inflection points were determined by minimising the residual sum of squares (defined as the observed values minus the population expectation at a given time point) on the transformed scale:

□ t_1_ = first inflection point: time point at which a treatment-induced change in tumour growth can be observed for an average mouse of a given patient line, and
□ t_2_ = second inflection point: time point at which a second (revertant) change in tumour growth (due to the end of treatment) could be observed for an average mouse of a given patient line, where 0 corresponds to the day of start of treatment for each PDX mouse.

Different model checks were performed to ensure that the selected model for each patient showed homoscedastic and normally distributed random effect predictions and residuals. Since no obvious violation of the model assumptions were noted, chosen models were taken forward and statistical inference results (p-values) trusted. P-values were subjected to multiplicity correction adjustments for within-patient analyses and comparisons.

### Collection and processing of dried blood spots

Blood spots were collected on day 1 (immediately before treatment start), 16 and 29 for PDX mice. Mice were immobilized in a stretcher/restrainer before ticking the tail with a needle. Upon squeezing the tail, ~50μl of blood were collected using a capillary lined with EDTA. The capillary was emptied into a 1.5ml microfuge tube and the blood was spotted onto Whatman FTA™ Classic Cards (Merck), and left to air dry for at least 15 minutes before storing at room temperature. For control experiments, blood spot samples were also collected from non-tumour bearing (healthy) NSG mice during terminal bleeds via cardiac puncture. Terminal bleeds were performed using syringes lined with EDTA, and 50μl of collected blood were subsequently spotted onto Whatman FTA™ Classic Cards. Again, cards were left to dry for 15 minutes.

In addition, dried blood spot samples were derived from 5 independent HGSOC patients (for use in dilution experiment; see **Supplementary Table 2** for patient information) by applying ~50μl of blood collected in K2-EDTA tubes to Whatman FTA™ Classic Cards.

### Shallow Whole Genome Sequencing (sWGS)

#### 1. Fresh frozen tumour tissue samples

Fresh frozen tissue pieces were homogenised using Soft tissue homogenizing CK14 tubes containing 1.4 mm ceramic beads (Bertin) on the Precellys tissue homogenizer instrument (Bertin). Lysates were subjected to DNA extraction using the AllPrep DNA/RNA Mini Kit (Qiagen) following manufacturer’s recommendations, and DNA was sheared to a fragment length of 200bp using the Covaris LE220 (120 sec at room temperature; 30% duty factor; 180W peak incident power; 50 cycles per burst).

Using the SMARTer Thruplex DNA-seq kit (Takara), 50ng of sheared DNA were prepared for sequencing following the recommended instructions with samples undergoing 5 PCR cycles for unique sample indexing and library amplification. Subsequently, AMPure XP beads were used (following manufacturer’s recommendations) to clean prepared libraries, which were then quantified and quality-checked using the Agilent 4200 TapeStation System (G2991AA). Pooled libraries were sequenced at low coverage on the HiSeq 4000 with single 50bp reads, at the CRUK CI Genomic Core Facility. Sequencing reads were aligned to the 1000 Genomes Project GRCh37-derived reference genome using the ‘BWA’ aligner (v.0.07.17) with default parameters.

#### 2. Dried blood spot samples

DNA from dried blood spots was extracted using the Qiagen Investigator kit (Qiagen) as previously described^26^ and eluted in 50μl elution buffer. High molecular weight genomic DNA (gDNA) was removed using right-side size selection with AMPure XP beads at a 1:1 and 7:1 bead:sample ratio (Beckman Coulter) described previously^26^, and eluted in 25μl water.

Before undergoing ThruPLEX Tag-seq library preparation (Takara), samples were concentrated to 10μl using a vacuum concentrator (SpeedVac). Samples were amplified for 14 to 16 cycles before undergoing the recommended bead clean up to remove remaining adapters. Quality control for library generation and quantification was done using a TapeStation (Agilent) before samples were submitted for sequencing on a NovaSeq 6000 SP (Illumina, paired-end 150bp), at the CRUK CI Genomic Core Facility.

### Analysis of dried blood spot sequencing data

Blood spot sequencing data was aligned to the human (hg19) and mouse genome (mm10) using |Xenomapper^27^. Reads overlapping with blacklisted regions for both human and mouse genomes were removed using the bedtools intersect function. Using Picard CollectInsertSizeMetrics, insert sizes were determined for the specific output files for each species. We computed a human ratio, that we call xenograft Tumour Fraction (xTF), for each sample by taking the total number of human reads >30bp fragment length and divided it by all reads (mouse and human) >30bp fragment length. Fragments below 30bp fragment length were excluded from the analysis as they tended to be noisy.

### Dilution series

To test the sensitivity and specificity of the human ratio metric, an *in silico* dilution experiment was performed using dried blood spot sequencing reads from five independent OV04 HGSOC patients (i.e. human reads only) and a healthy (non-tumour bearing) NSG mouse (i.e. mouse reads only). First, fastq files were aligned to the human (hg19) and mouse (mm10) reference genomes, respectively, to account for differences in sample quality, and to remove unmappable and duplicate reads. Resulting bam files were converted back to paired-end fastq files using the bedtools bamToFastq conversion utility. Mouse and human fastq files were then downsampled and merged to generate a seven-point dilution series containing 1%, 2%, 5%, 7%, 10%, 15% and 25% of human reads diluted in mouse reads for each of the 5 OV04 patients (35 samples in total). Paired-end fastq file pairs were then analysed with the Xenomapper pipeline, and human ratios (xTFs) estimated as described above. Estimated xTFs were then compared to expected human ratios based on *in silico* dilution mixtures.

### Absolute copy number analyses

We used the QDNAseq R package^29^ (v1.24.0) to count reads within 30 and 500kb bins, followed by read count correction for sequence mappability and GC content, and copy number segmentation. Resulting relative copy number data was then subjected to downstream analyses using the Rascal R package for ploidy and cellularity estimation and absolute copy number fitting as previously described^30^. For dried blood spot (DBS) samples, ploidy information from fitted tumour tissue samples from the same patient line were used to guide accurate ACN fitting. Note that DBS samples from healthy (non-tumour bearing) mice were automatically fitted to diploid ACN fits due to the absence of tumour reads and detectable somatic copy number aberrations (SCNAs).

Following ploidy and cellularity estimation, absolute copy number (ACN) profiles were generated for tumour tissues and DBS samples and subsequently correlated/compared across each 500kb bin. Putative driver amplifications were detected and identified using the Catalogue Of Somatic Mutations In Cancer (COSMIC; https://cancer.sanger.ac.uk/cosmic/help/cnv/overview) definitions and thresholds for high level amplifications and homozygous deletions: Gain: average genome ploidy ≤2.7 *and* total copy number ≥5; or average genome ploidy >2.7 *and* total copy number ≥9. Loss: average genome ploidy ≤2.7 *and* total copy number = 0; or average genome ploidy >2.7 *and* total copy number < (average genome ploidy - 2.7). Copy number signatures, as shown in **Supplementary Fig. 4**, were estimated as previously described^31^.

### Tagged-Amplicon Sequencing (TAm-Seq)

Small indels and single nucleotide variants were assessed across the coding regions of *TP53, BRCA1, BRCA2, MLH1, MSH2, MSH6, NF1, PMS2, PTEN, RAD51B, RAD51C, RAD51D*, and mutation hot spot regions for *BRAF, EGFR, KRAS*, and *PIK3CA* using the Tagged-Amplicon deep sequencing technology as previously reported^44^. Briefly, libraries were prepared in 48.48 Juno Access Array Integrated Fluidic Circuits chips (Fluidigm, PN 101-1926) on the IFC Controller AX instrument (Fluidigm), and libraries were sequenced by the CRUK CI Genomics Core Facility using 150bp paired-end mode on either the NovaSeq 6000 (SP flowcell) or HiSeq 4000 system. Reads were aligned to the GRCh37 reference genome using the ‘BWA-MEM’ aligner and variant calling was performed as previously described^45^.

### Haematoxylin and Eosin (H&E) and immunohistochemical p53 staining

H&E and immunohistochemical staining of p53 were carried out but the CRUK CI Histopathology Core Facility. H&E sections were stained following the Harris H&E staining protocol using a multi-stainer instrument (Leica ST5020). p53 staining was performed on 3μm FFPE sections using the Leica Bond Max fully automated IHC system. Antigen retrieval was performed using sodium citrate for 30 mins, and p53 was stained using the D07 Dako p53 antibody (1:1000).

### Data and code availability

Raw sequencing data from DBS samples will be uploaded to the European Genome-phenome Archive (EGA) database prior to publication. All supplementary data and all code required to reproduce the analyses and figures presented in this manuscript will be deposited on the Biostudies database.

## Acknowledgements

We would like to thank all patients who participated in and donated samples to this study. We thank Gemma Cronshaw and the biological resource unit (BRU) of the Cancer Research UK Cambridge Institute for their support with the animal models and performing weekly tumour measurement. We would like to thank the Cancer Research UK Cambridge Institute Genomics, IT & Scientific Computing and Histopathology core facilities for their support with various aspects of this work.

We also thank the Cancer Molecular Diagnostics Laboratory/Blood Processing Laboratory, which is supported by Cambridge NIHR Biomedical Research Centre, Cambridge Cancer Centre and the Mark Foundation of Cancer Research, who have performed blood collection, ctDNA isolation and bioinformatics analysis on the ovarian cancer patients. We acknowledge funding and support from Cancer Research UK, and the Cancer Research UK Cambridge Centre.

## Author Contributions

Conceptualization, C.M.S., K. Heider, N.R. and J.D.B.; Methodology, C.M.S., K.Heider, S.E.B., J.A.H., A.V. and M.V.; Software, C.M.S., K.Heider, and A.V.; Validation, C.M.S. and K.Heider; Formal Analysis, C.M.S., K.Heider, A.V. and D.L.C.; Investigation, C.M.S., K. Heider, J.B., S.E.B., J.A.H., A.A. and M.V.; Resources, C.M.S., K. Heider, J.B., J.A.H., S.E.B., A.A., A.V., D.L.C. and M.V.; Data Curation, C.M.S., K. Heider, J.B., and A.V.; Clinical Data, M.A.V.R. and K.Hosking; Writing - Original Draft, C.M.S., K. Heider, N.R. and J.D.B.; Writing - Review & Editing, C.M.S., K. Heider, D.L.C., A.A., M.A.V.R., M.V., N.R. and J.D.B.; Visualisation, C.M.S. and K. Heider; Supervision, C.M.S., K. Heider, M.V., N.R. and J.D.B.; Project Administration, C.M.S., K. Heider, M.V., N.R. and J.D.B.; Funding Acquisition, C.M.S., K.Heider, N.R. and J.D.B.

## Competing Financial Interests

Several of the authors are inventors and contributors on patents relating to methods for ctDNA analysis including methods described and used in this study. N.R. is an officer of Inivata Ltd. which commercialises ctDNA assays. J.D.B. is a founder of Tailor Bio. Both, Inivata and Tailor Bio, had no role in the conceptualization or design of the pre-clinical study, statistical analysis or decision to publish the manuscript.

## References

1. Deveson, I. W. et al. Evaluating the analytical validity of circulating tumor DNA sequencing assays for precision oncology. Nat. Biotechnol. 39, (2021).

2. Kilgour, E., Rothwell, D. G., Brady, G. & Dive, C. Liquid Biopsy-Based Biomarkers of Treatment Response and Resistance. Cancer Cell 37, 485–495 (2020).

3. Heitzer, E., Haque, I. S., Roberts, C. E. S. & Speicher, M. R. Current and future perspectives of liquid biopsies in genomics-driven oncology. Nat. Rev. Genet. 20, 71–88 (2019).

4. Rothwell, D. G. et al. Utility of ctDNA to support patient selection for early phase clinical trials: the TARGET study. Nat. Med. 25, 738–743 (2019).

5. Cohen, J. D. et al. Detection and localization of surgically resectable cancers with a multi-analyte blood test. Science (80-.). 359, 926–930 (2018).

6. Wan, J. C. M. et al. Liquid biopsies come of age: Towards implementation of circulating tumour DNA. Nat. Rev. Cancer 17, 223–238 (2017).

7. Cescon, D. W., Bratman, S. V., Chan, S. M. & Siu, L. L. Circulating tumor DNA and liquid biopsy in oncology. Nat. Cancer 1, 276–290 (2020).

8. Rolfo, C. et al. Liquid Biopsy for Advanced NSCLC: A Consensus Statement From the International Association for the Study of Lung Cancer. J. Thorac. Oncol. 16, 1647–1662 (2021).

9. Paracchini, L. et al. Genome-wide copy-number alterations in circulating tumor DNA as a novel biomarker for patients with high-grade serous ovarian cancer. Clin. Cancer Res. 27, 2549–2560 (2021).

10. Wan, J. C. M. et al. ctDNA monitoring using patient-specific sequencing and integration of variant reads. Sci. Transl. Med. 12, 1–17 (2020).

11. Zviran, A. et al. Genome-wide cell-free DNA mutational integration enables ultra-sensitive cancer monitoring. Nat. Med. 26, 1114–1124 (2020).

12. Abbosh, C. & Swanton, C. ctDNA: An emerging neoadjuvant biomarker in resectable solid tumors. PLOS Med. 18, e1003771 (2021).

13. Chen, M. & Zhao, H. Next-generation sequencing in liquid biopsy: cancer screening and early detection. Hum. Genomics 13, 34 (2019).

14. Adalsteinsson, V. A. et al. Scalable whole-exome sequencing of cell-free DNA reveals high concordance with metastatic tumors. Nat. Commun. 8, (2017).

15. Mouliere, F. et al. Enhanced detection of circulating tumor DNA by fragment size analysis. Sci. Transl. Med. 10, 1–14 (2018).

16. Ulz, P. et al. Inference of transcription factor binding from cell-free DNA enables tumor subtype prediction and early detection. Nat. Commun. 10, (2019).

17. Keller, L., Belloum, Y., Wikman, H. & Pantel, K. Clinical relevance of blood-based ctDNA analysis: mutation detection and beyond. Br. J. Cancer 124, 345–358 (2021).

18. Cristiano, S. et al. Genome-wide cell-free DNA fragmentation in patients with cancer. Nature 570, 385–389 (2019).

19. Markus, H. et al. Refined characterization of circulating tumor DNA through biological feature integration. medRxiv 1–13 (2021) doi:https://doi.org/10.1101/2021.08.11.21261907.

20. Ireson, C. R., Alavijeh, M. S., Palmer, A. M., Fowler, E. R. & Jones, H. J. The role of mouse tumour models in the discovery and development of anticancer drugs. Br. J. Cancer 121, 101–108 (2019).

21. Williams, J. A. Using pdx for preclinical cancer drug discovery: The evolving field. J. Clin. Med. 7, (2018).

22. Pearson, A. T. et al. Patient-derived xenograft (PDX) tumors increase growth rate with time. Oncotarget 7, 7993–8005 (2016).

23. Ice, R. J. et al. Drug responses are conserved across patient-derived xenograft models of melanoma leading to identification of novel drug combination therapies. Br. J. Cancer 122, 648–657 (2019).

24. Koessinger, A. L. et al. Quantitative in vivo bioluminescence imaging of orthotopic patient-derived glioblastoma xenografts. Sci. Rep. 10, 1–10 (2020).

25. Weissleder, R. Scaling down imaging: Molecular mapping of cancer in mice. Nat. Rev. Cancer 2, 11–18 (2002).

26. Heider, K. et al. Detection of ctDNA from Dried Blood Spots after DNA Size Selection. Clin. Chem. 66, 697–705 (2020).

27. Wakefield, M. J. Xenomapper: Mapping reads in a mixed species context. J. Open Source Softw. 1, 18 (2016).

28. Jahr, S. et al. DNA fragments in the blood plasma of cancer patients: Quantitations and evidence for their origin from apoptotic and necrotic cells. Cancer Res. 61, 1659–1665 (2001).

29. Scheinin, I. et al. DNA copy number analysis of fresh and formalin-fixed specimens by shallow whole-genome sequencing with identification and exclusion of problematic regions in the genome assembly. Genome Res. 24, 2022–2032 (2014).

30. Sauer, C. M. et al. Absolute copy number fitting from shallow whole genome sequencing data. bioRxiv 2021.07.19.452658 (2021).

31. Macintyre, G. et al. Copy number signatures and mutational processes in ovarian carcinoma. Nat. Genet. 50, 1262–1270 (2018).

32. O’Leary, B. et al. Early circulating tumor DNA dynamics and clonal selection with palbociclib and fulvestrant for breast cancer. Nat. Commun. 9, 1–10 (2018).

33. Chaisomchit, S., Wichajarn, R., Janejai, N. & Chareonsiriwatana, W. Stability of genomic DNA in dried blood spots stored on filter paper. Southeast Asian J. Trop. Med. Public Health 36, 270–273 (2005).

34. Lehmann-Werman, R. et al. Identification of tissue-specific cell death using methylation patterns of circulating DNA. Proc. Natl. Acad. Sci. U. S. A. 113, E1826–E1834 (2016).

35. Kurtz, D. M. et al. Enhanced detection of minimal residual disease by targeted sequencing of phased variants in circulating tumor DNA. Nat. Biotechnol. (2021) doi:10.1038/s41587-021-00981-w.

36. Abbosh, C. et al. Phylogenetic ctDNA analysis depicts early-stage lung cancer evolution. Nature 545, 446–451 (2017).

37. Parsons, H. A. et al. Sensitive Detection of Minimal Residual Disease in Patients Treated for Early-Stage Breast Cancer. Clin. Cancer Res. 26, 2556–2564 (2020).

38. Wickremsinhe, E. R. & Perkins, E. J. Using dried blood spot sampling to improve data quality and reduce animal use in mouse pharmacokinetic studies. J. Am. Assoc. Lab. Anim. Sci. 54, 139–144 (2015).

39. Prescott, M. J. & Lidster, K. Improving quality of science through better animal welfare: The NC3Rs strategy. Lab Anim. (NY). 46, 152–156 (2017).

40. Durschlag, M., Wurbel, H., Stauffacher, M. & Von Holst, D. Repeated blood collection in the laboratory mouse by tail incision-Modification of an old technique. Physiol. Behav. 60, 1565–1567 (1996).

41. Golde, W. T., Gollobin, P. & Rodriguez, L. L. A rapid, simple, and humane method for submandibular bleeding of mice using a lancet. Lab Anim. (NY). 34, 39–43 (2005).

42. Abatan, O. I., Welch, K. B. & Nemzek, J. A. Evaluation of saphenous venipuncture and modified tail-clip blood collection in mice. J. Am. Assoc. Lab. Anim. Sci. 47, 8–15 (2008).

43. Rago, C. et al. Serial assessment of human tumor burdens in mice by the analysis of circulating DNA. Cancer Res. 67, 9364–9370 (2007).

44. Forshew, T. et al. Noninvasive identification and monitoring of cancer mutations by targeted deep sequencing of plasma DNA. Sci. Transl. Med. 4, (2012).

45. Piskorz, A. M. et al. Methanol-based fixation is superior to buffered formalin for next-generation sequencing of DNA from clinical cancer samples. Ann. Oncol. 27, 532–539 (2016).

