## Supplementary Figures 1-6 for "Longitudinal monitoring of disease burden and response using ctDNA from dried blood spots in xenograft models"

### Supplementary Figures and Tables

***Supplementary Figure 1 – Examples of copy number profiles from DBS and tumour tissue samples for each of the four patient lines.***

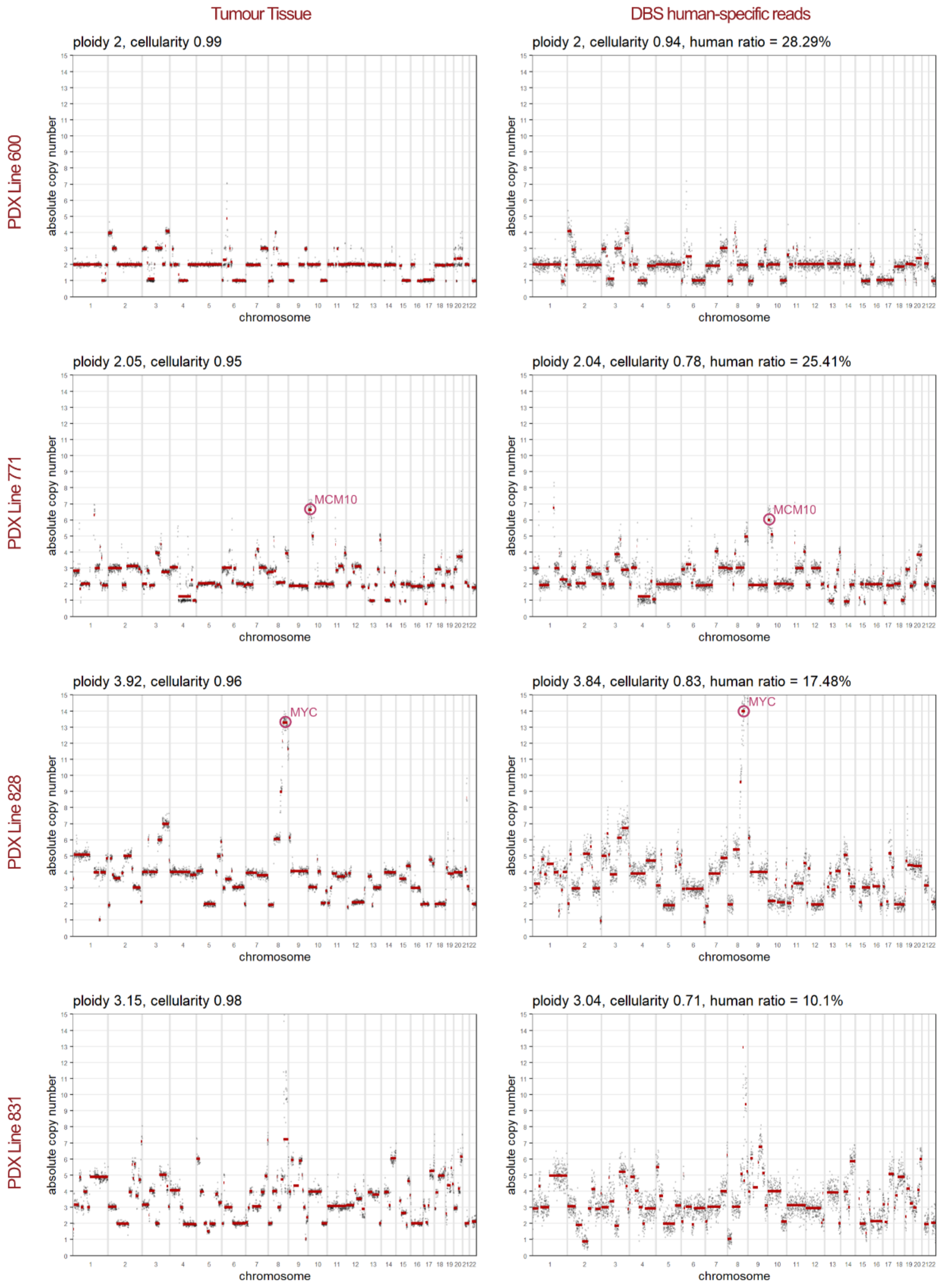


Comparison of absolute copy number (ACN) profiles obtained from sWGS data from a first generation PDX tumour tissues (left panel) and dried blood spot (DBS) samples from PDX mice (right panel) for patient 600, 771, 828, and patient 831. *MCM10* and *MYC* amplifications were detected in patients 771 and 828 samples, respectively, and are highlighted by pink circles.

***Supplementary Figure 2 – Correlation of SCNAs and detection of driver amplifications in DBS.***

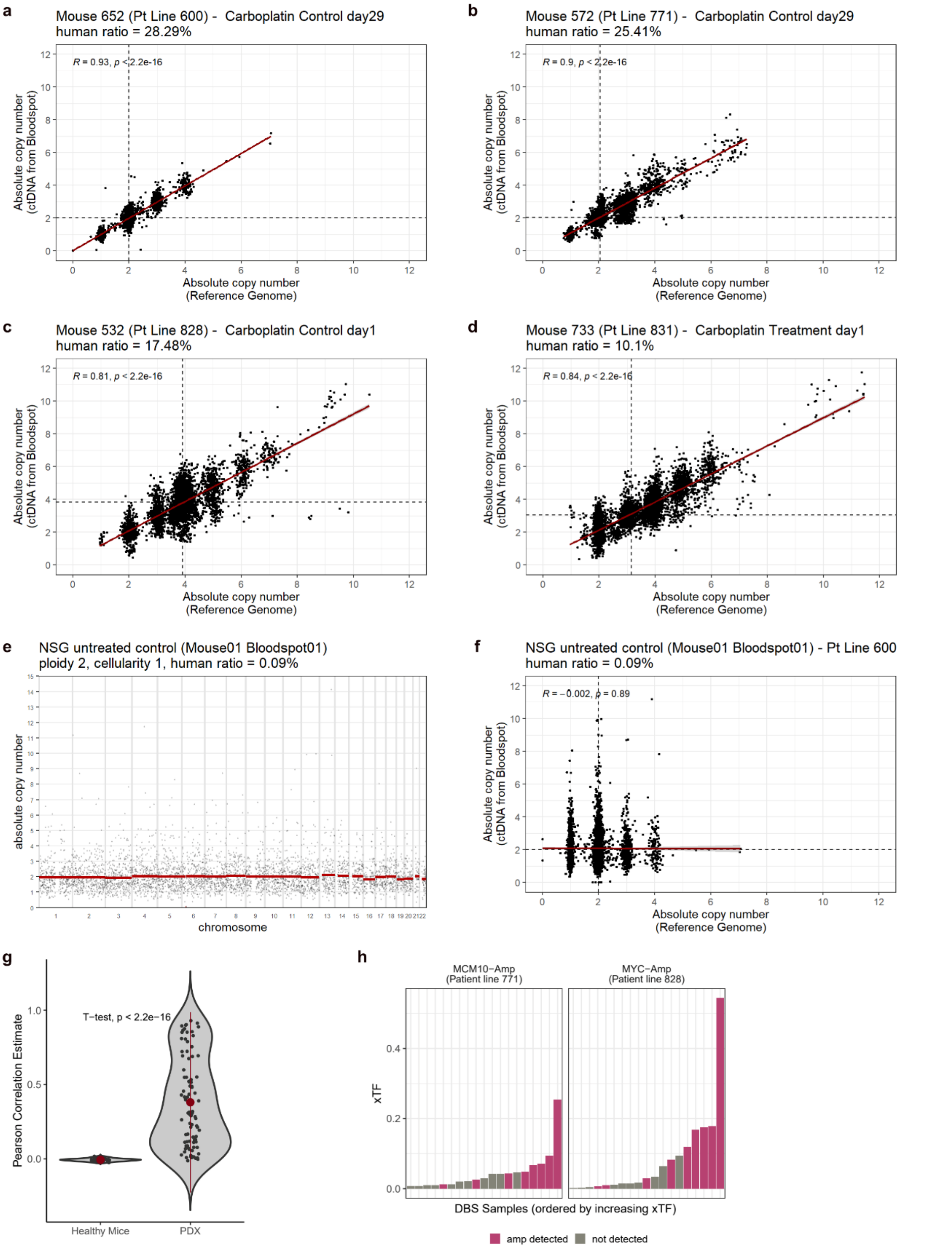


Pearson correlation plots comparing absolute copy number (ACN) profiles shown in **Supplementary Figure 1** for patient lines 600 (**a**), 771 (**b**), 828 (**c**) and 831 (**d**). Grey dashed lines indicated fitted ploidies for ACN profiles obtained from dried blood spots (horizontal line) and PDX tumour tissues (vertical line). (**e**) Example of an ACN profile obtained from a blood spot sample from a healthy non-tumour bearing mouse with an xTF of 0.0009 (0.09%). (**f**) Example correlation plot comparing the ACN profiles obtained from a blood spot from a healthy non-tumour bearing mouse and the first-generation tumour tissue from

PDX line 600. (**g**) Comparison of Pearson correlation estimates (correlating ACN profiles from blood spot and tumour tissue samples) between healthy (non-tumour bearing) and PDX mice (Welch t-test, $p < 2.2. x10^{-16}$; Wilcoxon test, $p < 2.2. x10^{-16}$). (**h**) Waterfall plot indicating samples for which putative driver amplifications were detected using COSMIC specifications (**see Methods**). Samples are arranged by increasing xTF values. Gene amplifications are detected in samples indicated in pink.

***Supplementary Figure 3 – Clinical treatment response, surgery outcome and time until first progression for the four pre-clinical study patients.***

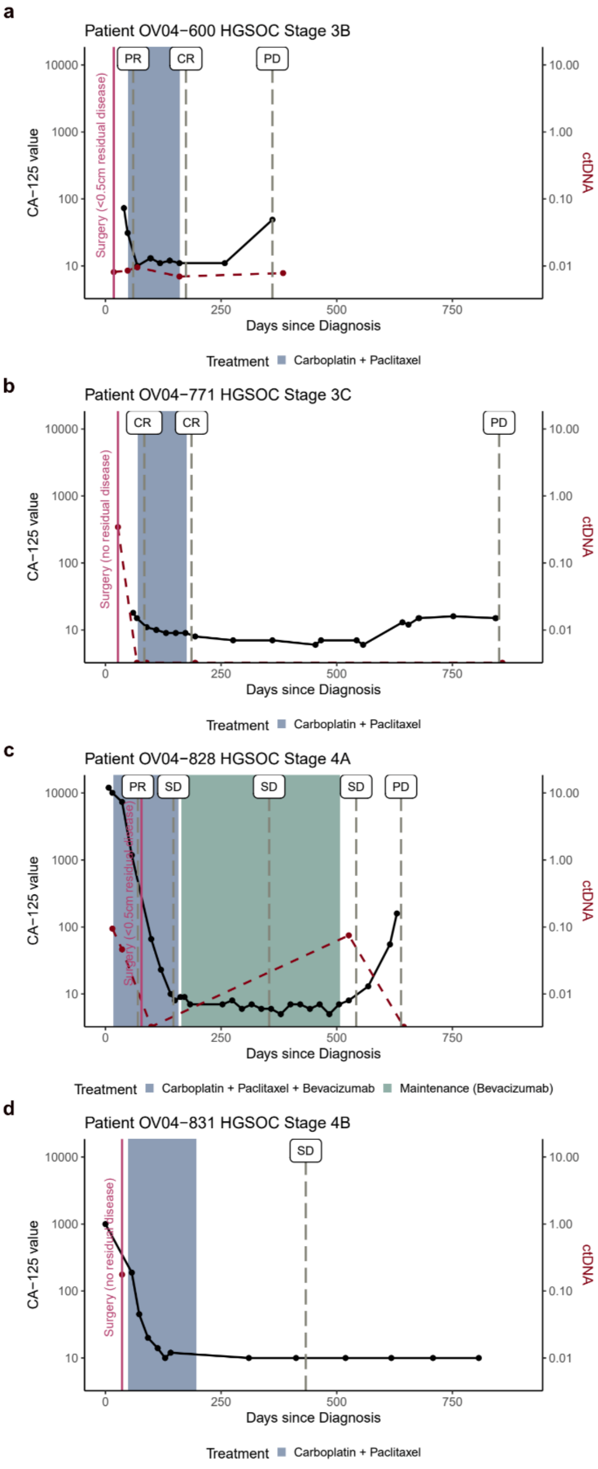


(**a-d**) CA-125 values (black line), treatment response assessments estimated via CT scans (vertical grey dashed lines), and ctDNA data, where available, (red dashed line), for HGSOC patients 600, 771, 828, and 831, respectively, over time. Surgery and additional treatment regimens are indicated by a pink vertical line and shaded boxes, respectively. (CR = Complete Response; PR = Partial Response; SD = Stable Disease; PD = Progressive Disease).

***Supplementary Figure 4 – Copy number signatures for each of the four patient lines.***

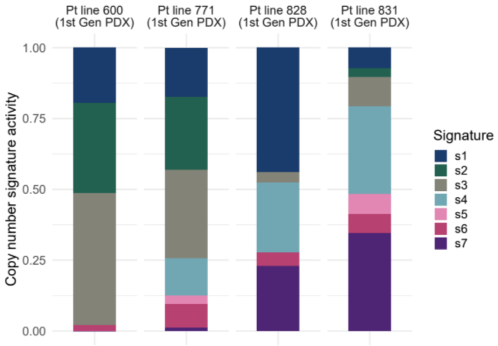


Stacked bar plots showing copy number signature activities for first generation PDX tissues derived from the four patients (patient 600, 771, 828 and 831) used in the pre-clinical HGSOC study.

***Supplementary Figure 5 – Histological features of patient and PDX tumour tissues.***

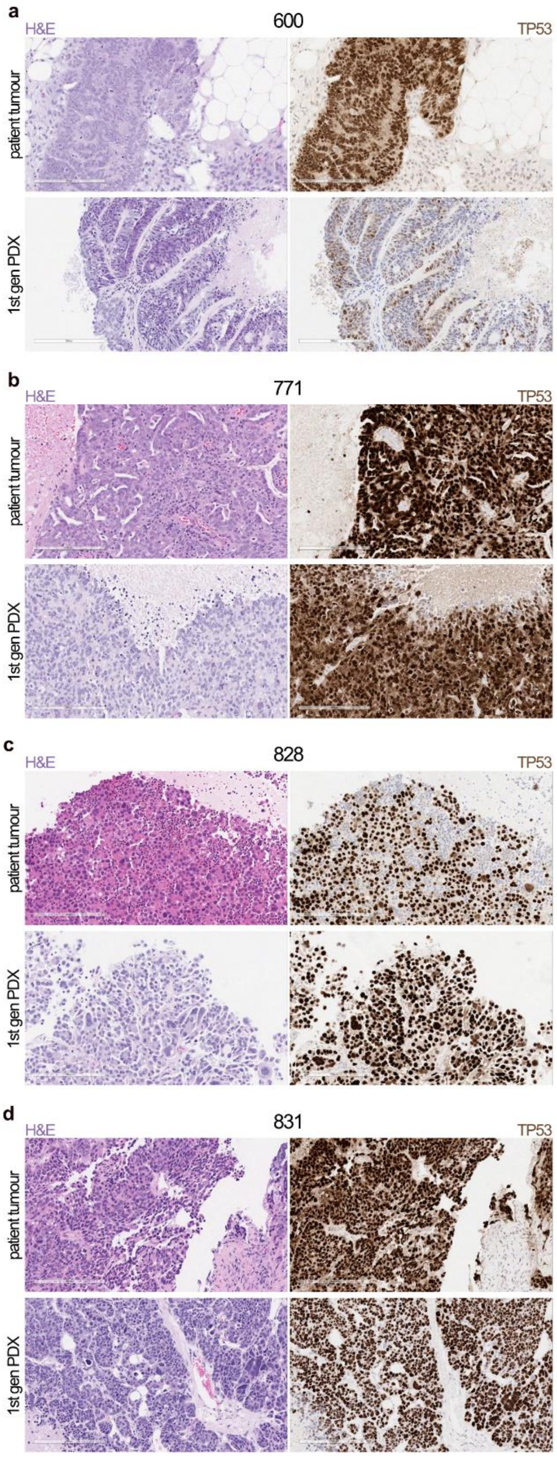


Comparison of haematoxylin and eosin (H&E, left panel) and *TP53* immunohistochemistry (IHC, right panel) stainings between patient and PDX tumour tissues for (**a**) patient 600, (**b**) patient 771, (**c**) patient 828, and (**d**) patient 831. The patient tumour is shown on the top, and the PDX tumour is shown on the bottom of each figure panel. Scale = 200µm.

***Supplementary Figure 6 – Molecular features of patient and PDX tumour tissues.***

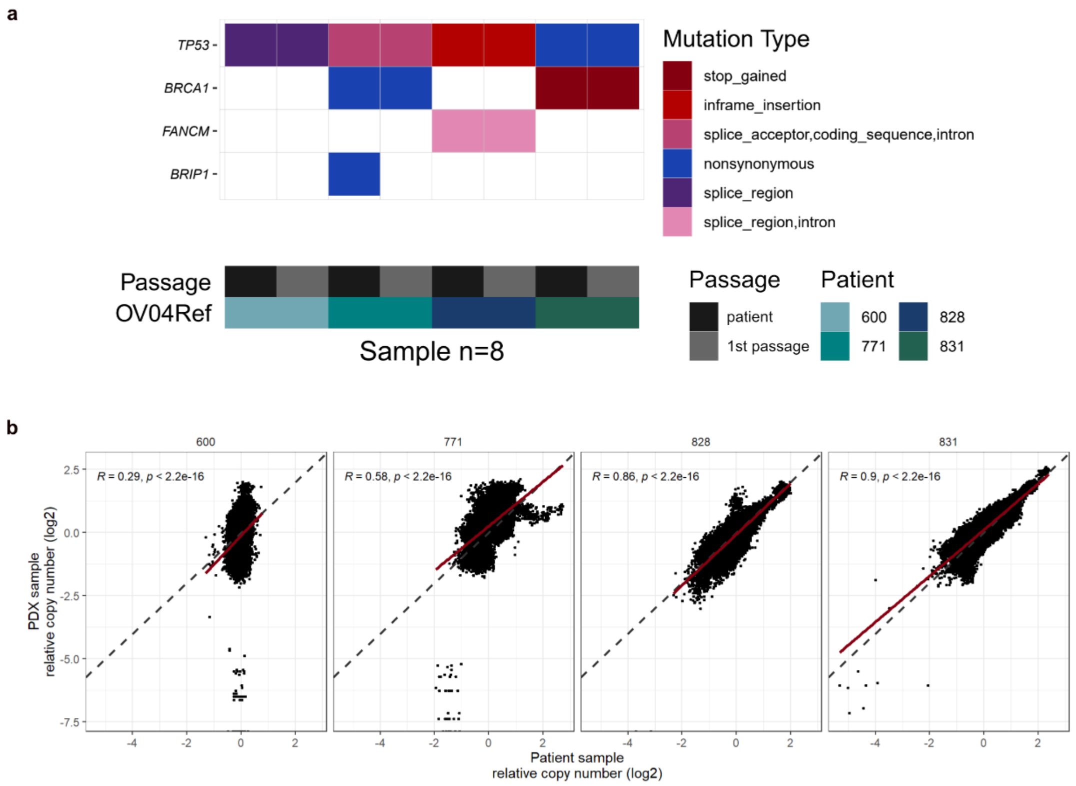


(**a**) Tagged-Amplicon Sequencing (TAm-Seq) mutational data comparing patient tumour tissues, and first generation PDX tumour tissues for each of the four patient lines, as indicated by different shades of blue/green. Patient tissues are indicated by dark grey blocks, whereas corresponding PDX tumour tissues are indicated by light grey blocks. (**b**) Comparison of relative copy number data (log2 scale) obtained from shallow whole genome sequencing between the original patient tumour tissue, and the first generation PDX tumour tissue for each of the four patient lines. Diagonal grey dashed line indicates a slope of 1. Note that the original tumour tissue for patient 600 had very low tumour purity (<10%) resulting in a mostly flat copy number profile and consequently a poorer correlation when compared to the associated pure PDX tumour tissue.

***Supplementary Figure 7 – Overview of pre-clinical patients and PDX lines.***

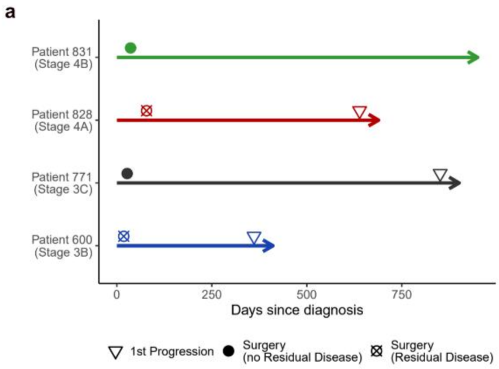


**b**

| PATIENT | | | | | | | | | | PDX MODELS |
|---|---|---|---|---|---|---|---|---|---|---|
| **ID** | **Stage** | **Age at diagnosis** | **Time to relapse** | **Overall survival** | ***TP53*** | ***BRCA1/2*** | **other** | **Duration treatment effect** | **Time to regrowth** | **Time to endpoint (T/C)** |
| **600** | IIIB | 46 | 361 days | 1322 days | c.994-1G>C | - | - | 22 days | 23 days | 67.6 / 39.4 days |
| **771** | IIIC | 51 | 851 days | - | c.783-5_785del | - | ER+ | 28 days | 33 days | 62.0 / 31.6 days |
| **828** | IVA | 60 | 639 days | 1662 days | c.823_825dup | sBRCA1 translocation | *MYC* Amp | 41 days | 44 days | 151.3 / 28 days |
| **831** | IVB | 61 | - | - | c.584T>C | gBRCA1m | ER+ | 39 days | 41 days | 67.8 / 35.0 days |

(**a**) Overview of clinical timelines until first disease progression in the four HGSOC patients. (**b**) Table summarising clinical, genomic and PDX tumour growth kinetics information for all four patients included in this study. Patient 600, 771, and 831 underwent primary surgery prior to treatment, whereas patient 828 received neoadjuvant treatment prior to interval debulking surgery and further treatment. Both patient 600 and 828 had residual disease following surgery.

***Supplementary Table 1 - Tumour growth modelling***

The following Table shows the unadjusted ('raw' columns) and adjusted ('adj' columns) p-values and star-based significance levels for each patient line corresponding to the following comparions:

**Contrast 1**: Difference in tumour volume between the control and treatment groups at treatment start
**Contrast 2**: Difference in tumour growth between the control and treatment groups before time inflection point $t_1$ (i.e. before start of treatment effect)
**Contrast 3**: Difference in tumour growth before and after time inflection point $t_1$ for treated mice (a significant effect would correspond to a significant tumour growth change due to treatment)
**Contrast 4**: Difference in tumour growth before and after inflection point $t_2$ for treated mice (a significant effect would correspond to a significant tumour growth change due to end of treatment)
**Contrast 5**: Difference in tumour growth before inflection point $t_1$ and after inflection point $t_2$ for treated mice (a significant effect would correspond to a significant tumour growth change before and after treatment effect)
**Contrast 6**: Difference in tumour growth between the control and treatment groups after inflection point $t_2$ (where the tumour growth is assumed constant through time for mice of the control group)

**Patient line 600**

| contrasts | pval.raw | sig.raw | pval.adj | sig.adj |
|---|---|---|---|---|
| Contrasts1 | 0.700861 | | 1 | |
| Contrasts2 | 0.9252195 | | 1 | |
| Contrasts3 | 1.88528E-05 | *** | 0.000364298 | *** |
| Contrasts4 | 3.53503E-10 | *** | 5.6196E-09 | *** |
| Contrasts5 | 0.8592234 | | 1 | |
| Contrasts6 | 0.5726736 | | 1 | |

**Patient line 771**

| contrasts | pval.raw | sig.raw | pval.adj | sig.adj |
|---|---|---|---|---|
| Contrasts1 | 0.2710158 | | 1 | |
| Contrasts2 | 0.4120366 | | 1 | |
| Contrasts3 | 1.22636E-09 | *** | 1.53545E-08 | *** |
| Contrasts4 | 3.43222E-08 | *** | 4.72356E-07 | *** |
| Contrasts5 | 0.1996257 | | 1 | |
| Contrasts6 | 0.02762937 | * | 0.4921333 | |

**Patient line 828**

| contrasts | pval.raw | sig.raw | pval.adj | sig.adj |
|---|---|---|---|---|
| Contrasts1 | 0.2054286 | | 1 | |
| Contrasts2 | 0.5641273 | | 1 | |
| Contrasts3 | 6.66134E-16 | *** | 1.5099E-14 | *** |
| Contrasts4 | 0 | *** | 0 | *** |
| Contrasts5 | 0.2650988 | | 1 | |
| Contrasts6 | 0.3385248 | | 1 | |

**Patient line 831**

| contrasts | pval.raw | sig.raw | pval.adj | sig.adj |
|---|---|---|---|---|
| Contrasts1 | 0.9144219 | | 1 | |
| Contrasts2 | 0.1758687 | | 1 | |
| Contrasts3 | 1.51909E-10 | *** | 2.21377E-09 | *** |
| Contrasts4 | 6.17251E-08 | *** | 1.14541E-06 | *** |
| Contrasts5 | 0.00387063 | ** | 0.07970715 | . |
| Contrasts6 | 0.09430599 | . | 1 | |

***Supplementary Table 2 – Patient overview for DBS dilution series analysis***

| Patient | StudyID | Histology | Stage | Barcode | SequencingID |
|---|---:|---|---|---|---|
| Patient 1 | 1132 | HGSOC | IVB | D708tp-D501tp | SLX-10615 |
| Patient 2 | 461 | HGSOC | IIIC | D708tp-D502tp | SLX-10615 |
| Patient 3 | 1117 | HGSOC | IC | D708tp-D503tp | SLX-10615 |
| Patient 4 | 1020 | HGSOC | IIIC | D708tp-D504tp | SLX-10615 |
| Patient 5 | 628 | HGSOC | IV | D708tp-D505tp | SLX-10615 |
